# Human neuronal firing is modulated by the frequency of local field potential oscillations

**DOI:** 10.1101/2025.09.03.674012

**Authors:** Zahra Jourahmad, Raissa K. Mathura, Layth S. Mattar, Melissa C. Franch, Danika L. Paulo, Mohammed Hasen, Nicole R. Provenza, Benjamin Y. Hayden, Sameer A. Sheth, Eleonora Bartoli, Andrew J. Watrous

## Abstract

Neural oscillations play a critical role in shaping neuronal firing patterns. While phase-locked neuronal firing (“phase tuning”) has been extensively studied in animal models and human invasive recordings, much less is known about whether neurons show preferential firing at specific oscillatory frequencies, termed frequency tuning. Here, we employ human intracranial recordings across several brain regions including hippocampus, entorhinal cortex, anterior and posterior cingulate cortex, and orbitofrontal cortex to test the hypothesis that neurons exhibit frequency-specific firing. We analyzed 357 single units recorded simultaneously with local field potentials in 19 neurosurgical patients during awake resting. We estimated the instantaneous frequency of the LFP using adaptive spectral decomposition and assessed frequency tuning of each neuron while controlling for changes in firing rate unrelated to frequency changes. We found 27% neurons exhibited increased or decreased firing within specific frequencies, most commonly within the low-theta range (<10 Hz). Neurons exhibiting frequency tuning were distinct from those displaying phase tuning, and both types of tuning were observed across multiple brain regions with no anatomical preference. Together, our results demonstrate that the instantaneous frequency of neural oscillations modulates neuronal firing which may serve as an additional mechanism for information processing in the human brain, opening new avenues for frequency-targeted neural stimulation.

## Introduction

Neuronal spiking activity is the fundamental unit of information processing in the brain, traditionally understood to encode information through firing rates (1). Additionally, research in both human (2–7) and non-human (8) models suggests that neuronal firing relative to local field potential (LFP) oscillations provide complementary mechanisms for neural coding. LFPs, which reflect the summed synaptic activity of local neuronal populations as well as non-synaptic currents, offer insight into the underlying dynamics of neural circuits (9–11). These fluctuations represent large-scale network activity that can influence the timing of neuronal firing (8,12) and may facilitate or suppress spiking depending on the phase and frequency of oscillatory activity (13). A growing body of research has shown that fluctuations in network-level oscillations are systematically related to single-neuron spiking activity (12). This interaction is thought to help structure neural activity in ways that support higher-order cognitive functions, such as memory and decision-making, by coordinating information flow across distributed brain regions (8,14,15).

Generally, oscillatory signals can be decomposed into an amplitude, phase, and frequency component, and, in principle, each of these aspects of the neural signal may modulate neuronal firing. Regarding phase, studies have highlighted the importance of phase coding, where the precise timing of neuronal firing relative to the phase of ongoing oscillations conveys behaviorally relevant information (16). While early studies focused on phase coding via phase precession (17) in rodent hippocampus (a relatively rhythmic, regular oscillatory environment), recent findings suggest that phase coding can also occur in highly irregular oscillatory environments, as observed in bats (18) and humans (5,19). This phase coding provides a structured temporal scaffold that can regulate when neurons are most excitable (20). For example, recent evidence suggests that neural activity tends to occur at specific phases of low-frequency oscillations, and that this phase preference is predictive of sensory processing, memory encoding, and retrieval success (6,7,15,21). These findings suggest that the phase alignment helps organize neural activity into content-specific patterns, offering a temporally precise mechanism for representing and accessing information in the brain. Notably, Qasim et al. demonstrated phase precession in human hippocampal and entorhinal neurons spanning a low-frequency range (2-10 Hz) extending beyond the narrower theta range typically described in rodents (19). They proposed that this flexible phase advancement may reflect a frequency-based coding scheme, in which information is encoded not only by phase, but also by the relative relationship between spike rhythmicity and the instantaneous LFP frequency. Extending this view, Schonhaut et al. showed that hippocampal oscillations synchronize neuronal firing both locally and in connected medial temporal regions, with neurons in different areas preferentially locking to distinct hippocampal frequency bands. Based on these findings, we hypothesized that slow oscillatory frequencies are not merely an “idling” rhythm but instead actively influence neuronal firing (6).

Prior modeling work by Cohen introduced the concept of frequency sliding, the dynamic, moment-to-moment fluctuations in the peak frequency of brain oscillations, and demonstrated that these shifts can influence spike timing and firing thresholds in biophysically realistic neurons (13). These subtle changes in frequency were proposed to reflect variations in input strength and network excitability, thereby modulating neuronal responsiveness to synaptic input. Complementing this view, experimental studies have shown that neurons possess intrinsic frequency preferences –a phenomenon known as resonance– where subthreshold membrane potentials are selectively amplified by inputs at specific frequencies (22). Oscillatory activity in the brain is thought to play a critical role in organizing neural computation by temporally coordinating spiking activity across distributed circuits. For example, the Spectro-Contextual Encoding and Retrieval Theory posits that frequency-specific oscillatory patterns serve as a contextual framework for binding and reactivating distributed representations during memory processes (23). At the core of this theory is the idea that frequency selectivity enables neurons to be differentially engaged by distinct oscillatory inputs based on their biophysical properties. Building on these theoretical and empirical foundations, it remains an open question whether such frequency-selective spiking responses can be directly observed using human single neuron recordings *in vivo*.

To elucidate how local oscillatory dynamics shape neuronal firing patterns, we simultaneously recorded LFPs and single-unit activity (SUA) in patients with epilepsy undergoing intracranial EEG monitoring for seizure localization. Motivated by Cohen’s frequency-sliding framework, we hypothesized that instantaneous oscillatory frequency may modulate neuronal firing. We asked whether human single neuron firing changes as a function of the instantaneous oscillatory frequencies in the LFP, examining this relationship across a wide range of frequencies and brain regions.

## Results

We analyzed neural data from a total of 19 patients (**Supplementary Table 1**) with pharmacoresistant epilepsy during resting state. Each patient was implanted with 2-7 Behnke-Fried probes, with locations spanning the hippocampus (Hipp), entorhinal cortex (EC), anterior cingulate cortex (ACC), posterior cingulate cortex (PCC), and orbitofrontal cortex (OFC) (**Figure 1A**). LFPs were obtained from the macroelectrode closest to the bundle of microwires used to isolate SUA. In total, the signal from 84 macroelectrodes was used for the LFP analysis (**Table 1**). SUA was recorded from microwires (n=8) in proximity of each macroelectrode (**Figure 1B**). The number of well-isolated single-units per microwire varied, typically ranging from zero to three, though in rare instances up to four distinct neurons were detected on a single wire. In total, 357 single-units were included in the analysis (**Table 1**).

**Figure 1.**
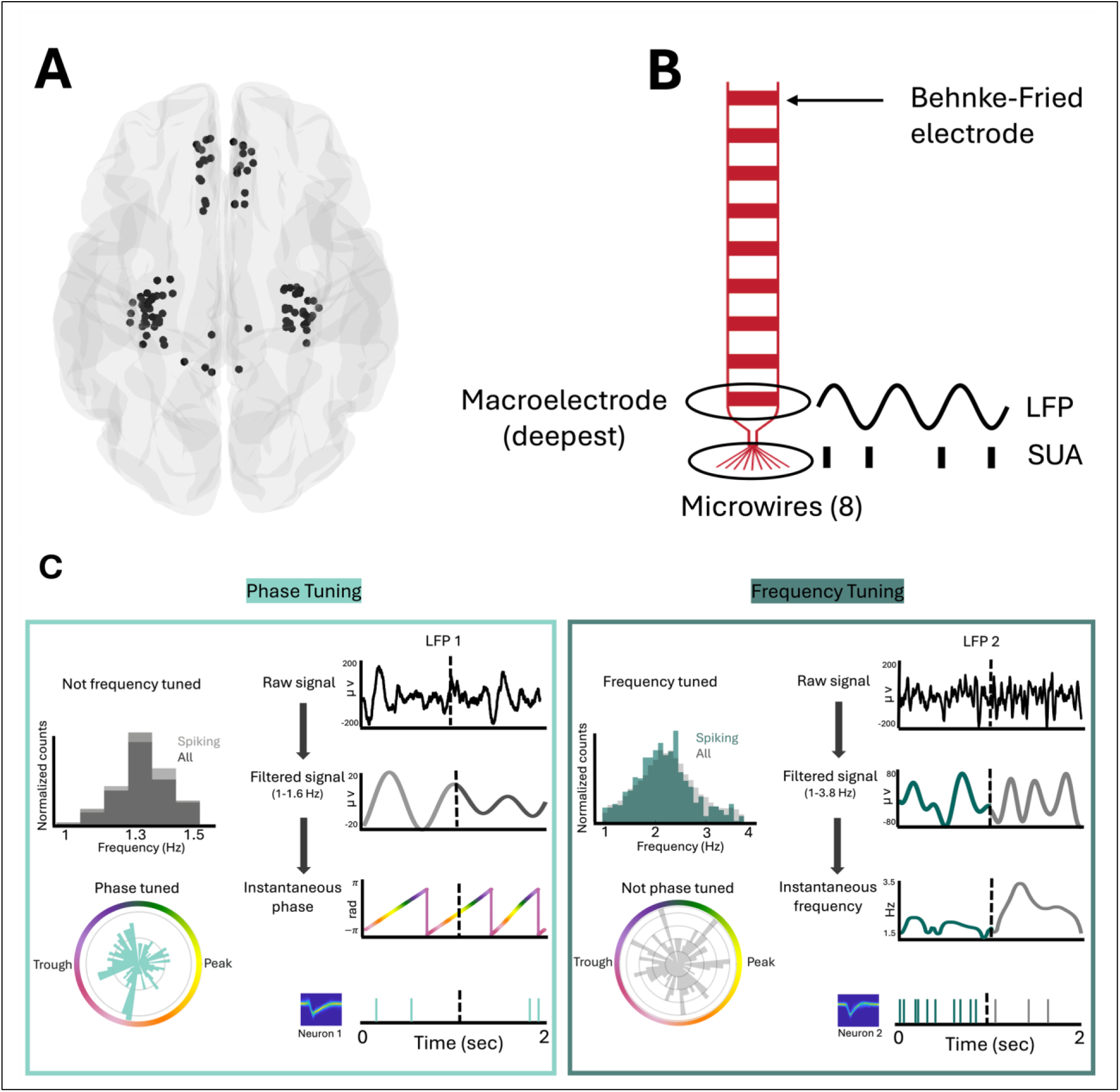
Anatomical recording sites and analytical framework for frequency and phase tuning. **A.** Anatomical location of the deepest macroelectrode contact for each patient (n = 84) represented in MNI space using the *fs average* brain template. Each point represents a macroelectrode used for LFP recording. **B.** Schematic of one representative Behnke-Fried electrode. LFPs were analyzed from the macroelectrode closest to the tip of the probe, and SUA was extracted from the 8 adjacent microwires. **C.** Analytical framework for testing frequency and phase-tuning hypotheses with examples. The raw LFP signal is decomposed into its instantaneous frequencies and phases, and spike times are aligned to these features. Frequency and phase tuning are assessed using the KS test and the Rayleigh test, respectively. The left panel shows phase tuning in a different neuron, which aligns its spiking activity to the trough of the oscillation within the first decomposed LFP band. This neuron does not display frequency tuning. In contrast, the right panel illustrates frequency tuning in a neuron relative to the first decomposed band of the LFP. Although the dominant frequency in the signal is 2.5 Hz, the neuron exhibits increased spiking density at 2 Hz. Notably, this neuron does not exhibit phase tuning.

**Table 1.** Summary of single-unit recording coverage by region.

| Region | #Patients | #Macros | #Single Units | Firing rate (95% CI) |
| --- | --- | --- | --- | --- |
| Hipp | 17 | 38 | 147 | 2.63 (2.10, 3.20) |
| EC | 6 | 12 | 70 | 2.5 (1.95, 3.10) |
| ACC | 14 | 25 | 100 | 2.9 (2.45, 3.47) |
| PCC | 5 | 5 | 17 | 3.03 (1.38, 4.68) |
| OFC | 3 | 4 | 23 | 0.83 (0.70, 0.97) |

We examined whether neuronal firing is influenced by the instantaneous phase or frequency of the LFP, as exemplified in **Figure 1C**, showcasing two representative neurons and our analytical framework to identify phase and frequency tuning.

Prior evidence in humans suggests variable theta frequencies across brain regions (2,19,24) and patients (6). We therefore used a previously validated adaptive spectral decomposition algorithm (25) to accommodate variable spectral content (**Supplementary Figure 1**) and to yield optimized phase and frequency estimates for the LFP signals (**Figure 2A,B**) compared to using classical frequency bands (**Figure 2C,D**).

**Figure 2.**
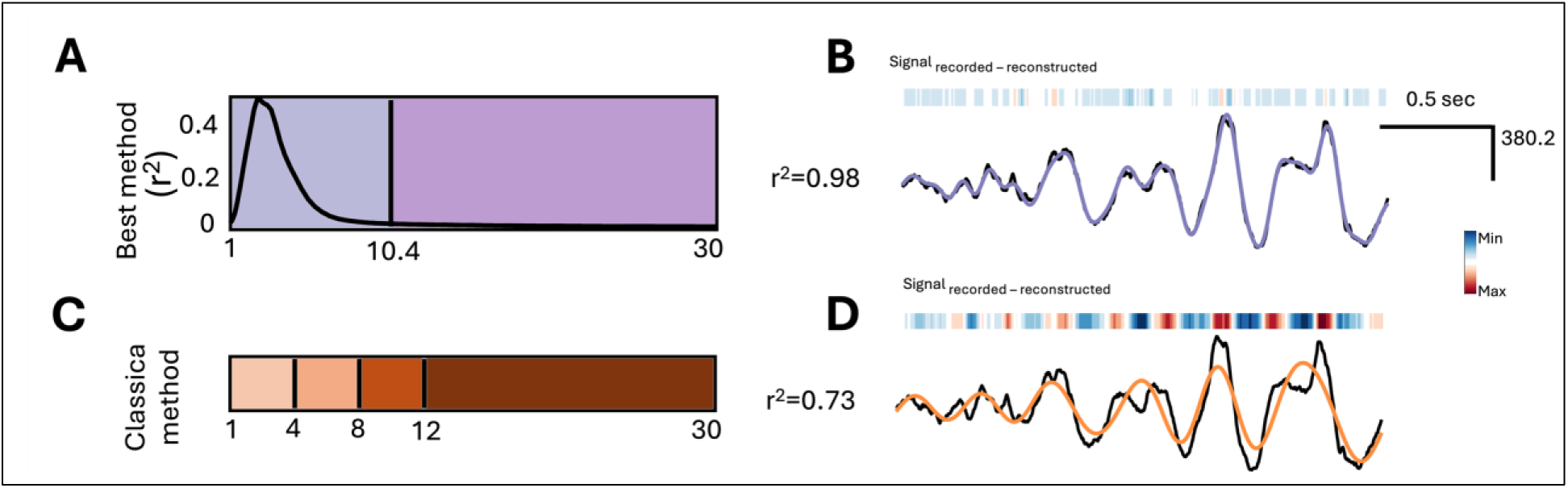
Comparison of signal decomposition methods and their effectiveness in reconstructing neural signals. **A.** Example of signal decomposition using the ORCA-selected method, in which the signal is decomposed into two data-driven frequency bands. The x-axis represents frequency, and the y-axis shows the explained variance attributable to each frequency component. As depicted, the maximum explained variance (∼0.5) occurs at ∼3 Hz. **B.** A 2-second segment showing the raw LFP (black) overlaid with the reconstructed signal (purple) using the ORCA-derived bands. The high goodness-of-fit (0.98) indicates excellent reconstruction. **C.** Example of signal decomposition using classical methods, dividing the same signal into four canonical frequency bands. **D.** A 2-second segment showing the raw LFP (black) overlaid with the reconstructed signal (orange) using the classical bands. The reconstruction remains lower than ORCA (0.73).

### A Subset of Human Single Neurons Displays Phase Tuning

Our phase tuning analyses revealed that 15% of neurons (n = 54/357) displayed significant phase tuning (binomial test, chance level = 0.01, p < 0.0001). For neurons showing significant phase tuning, we further examined the specific phases of the oscillatory cycle that contributed to this effect. We extracted the mean preferred phase of spiking from the circular distribution of spike phases for each neuron with significant phase locking, as determined by the Rayleigh test. This analysis revealed that phase preferences were not uniformly distributed but instead clustered around distinct phases of the cycle. Overall, the majority of neurons exhibited a unimodal phase preference. Specifically, 17 neurons exhibited phase preference near the peak of the oscillation (around 0 radians), 33 neurons preferred phases near the trough (around π), and 4 neurons showed tuning around intermediate phases, closer to π /2 or -π/2 positions between peak and trough on the cycle. **Figure 3A** shows an example phase tuned neuron whose spike times are displayed above a 2 second LFP trace; spikes occurred predominantly near the peak of the oscillation. We assessed whether this neuron exhibited frequency tuning by comparing instantaneous frequencies during spiking to those across the entire recording using the Kolmogorov–Smirnov (KS) test (**Figure 3B–D**).To quantify phase-related modulation, we applied the Rayleigh test, which revealed that spikes were non-uniformly distributed across phases, with a clear preference for the peak (p <0.001; **Figure 3E**).

**Figure 3.**
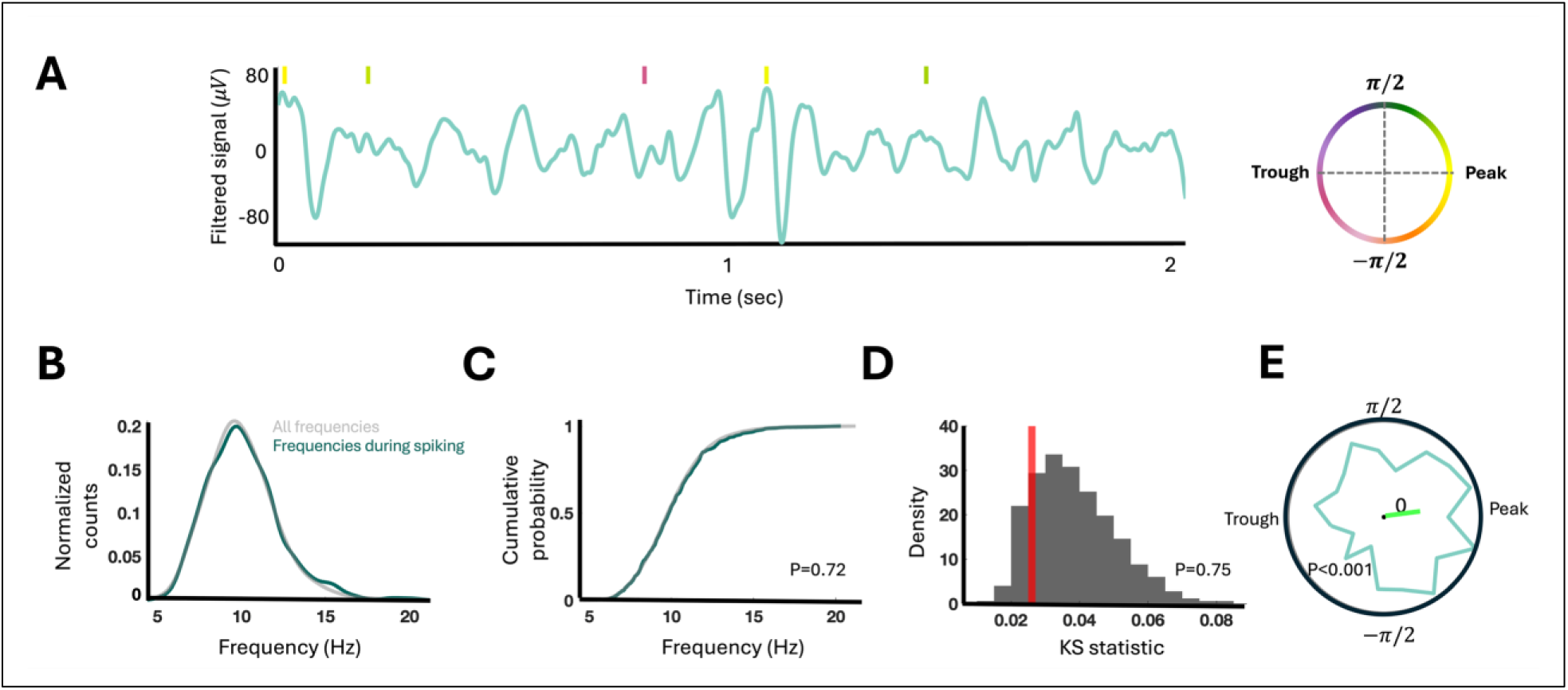
Example of phase tuning in a single neuron. **A.** A 2-second segment of the filtered LFP signal, with spike times shown as vertical ticks. Tick colors correspond to the instantaneous phase at the time of the spike, as indicated by the adjacent colorbar: pink/purple tones represent trough phases, while yellow/orange tones indicate peak phases. Spikes are preferentially aligned to the peak of the oscillation, indicating phase tuning. **B.** Probability distributions of instantaneous frequency during spike times (pine green) and across all times (gray), indicating no appreciable shift in frequency distribution during spiking. **C.** Cumulative distributions of instantaneous frequencies at spike times and across all times were compared using the KS test, revealing no significant difference. **D.** Permutation-based null distribution of the maximum cumulative difference (D-statistic), with the observed value (red line) falling within the null distribution, confirming the absence of statistically significant frequency tuning **E.** Circular phase distribution and corresponding mean vector (neon green), demonstrating significant phase tuning aligned with the peak of the oscillation.

To account for the potential influence of non-sinusoidal oscillations on phase tuning, we compared the distribution of phases for phase-tuned cells to the overall phase distribution of the LFP. In our analysis, 94% of phase tuned neurons exhibited a phase preference that significantly differed from the global LFP phase distribution (p < 0.05, Watson-Williams test). This finding indicates that phase-tuned activity is unlikely to be an artifact of non-sinusoidal waveform distortions.

Next, to determine whether specific phase preferences dominated within brain regions, we performed region-specific Rayleigh tests. A statistically significant phase preference was found in the ACC, where spikes were preferentially locked to the trough of the oscillation (Rayleigh test, p = 0.002). In contrast, the Hipp (p = 0.1) and EC (p = 0.2) did not show statistically significant clustering, although their mean preferred phases were also aligned with the descending phase (**Supplementary Figure 2**). Spiking near the trough is consistent with previous findings and may reflect increased neuronal excitability during this phase of the cycle, which facilitates neuronal firing (2,26,27).

### A Different Subset of Human Single Neurons Displays Frequency Tuning

We next tested neurons for frequency tuning. **Figure 4A** illustrates a neuron showing frequency tuning; spikes occurred predominantly during the low-frequency portion of the 2 second LFP, relative to the dominant oscillatory frequency (∼4 Hz).The KS test confirmed a significant difference between the frequency distribution at spike times and the overall distribution, and this effect persisted when compared to a permutation-based null distribution of the KS statistic (p <0.001; **Figure 4B–D**). In this neuron, the Rayleigh test indicated no significant phase preference, and the circular histogram of spike phases showed no clear clustering (**Figure 4E**). Our analysis revealed that 8% of all neurons (n = 27/357) exhibited significant frequency-tuning (binomial test, chance level = 0.01, p < 0.0001).

**Figure 4.**
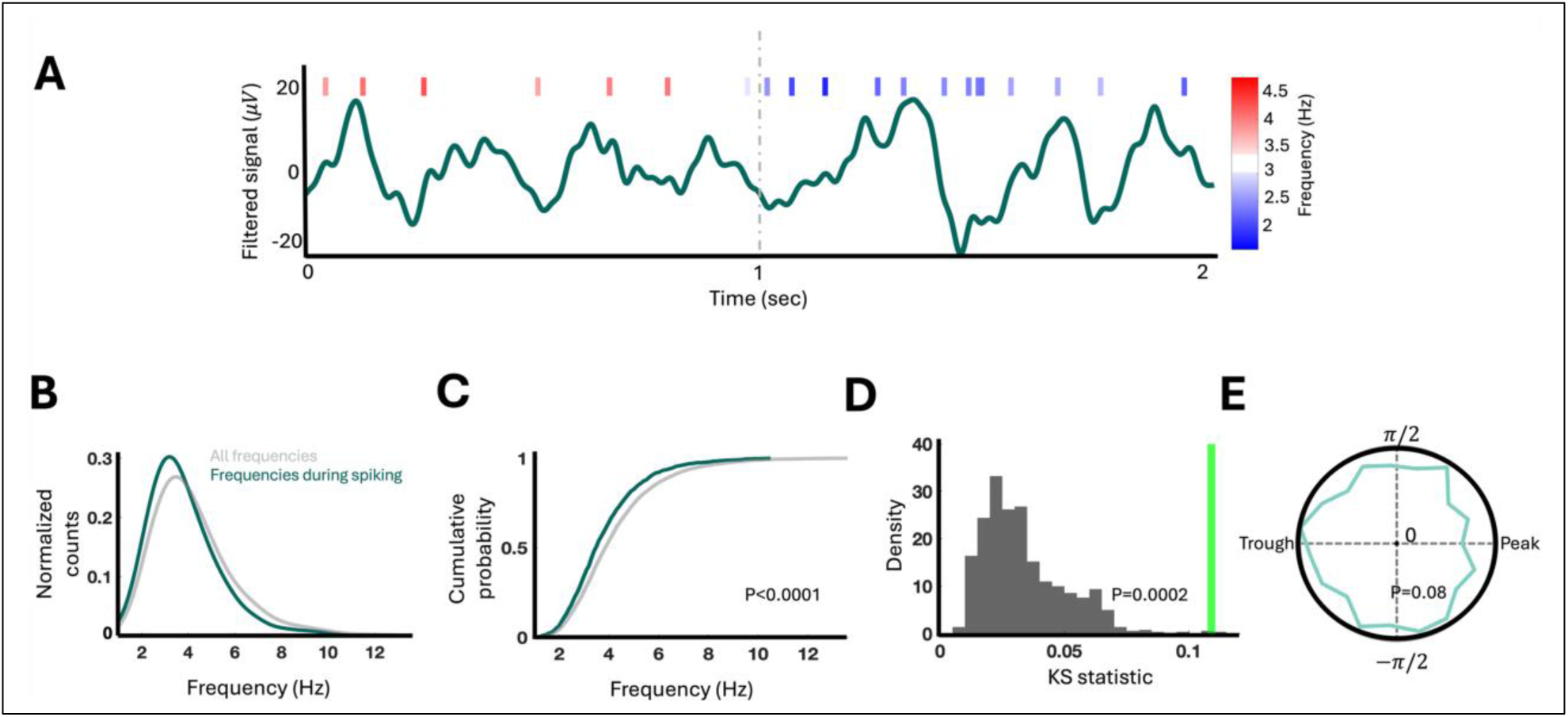
Example of frequency tuning in a single neuron. **A.** A 2-second segment of the filtered LFP signal, with spike times shown as vertical ticks. Spikes are color-coded: blue for those occurring during lower-frequency epochs (∼3 Hz) and red for those during higher-frequency epochs (∼4 Hz), illustrating a preference for spikes to occur during lower oscillation. As the frequency of the signal transitions from ∼4 Hz to ∼3 Hz, the spike rates increase, reflecting enhanced neuronal firing during slower oscillatory periods. **B.** Probability distributions of instantaneous frequency during spike times (pine green) and all times (gray), illustrating a shift in instantaneous frequencies during spiking compared to all frequencies. **C.** Cumulative distributions of instantaneous frequencies at spike times and across all times were compared using the KS test, revealing a significant difference. **D.** Permutation-based null distribution of the maximum cumulative difference (D-statistic), with the observed value (neon green line) falling outside the null distribution, confirming statistically significant frequency tuning. **E.** Circular phase distribution from the same neuron, showing no evidence of phase tuning.

For neurons showing frequency tuning, we further identified the specific frequency ranges contributing to this effect by comparing the probability density of instantaneous frequencies at spike times to all times. To do so we computed the difference in probability densities at each frequency and assessed its significance by comparing the observed difference to a null distribution generated through permutation (see Methods). This analysis revealed three distinct patterns of frequency-related modulation (**Figure 5**). The most common pattern was a bidirectional modulation observed in 18 neurons (67% of the neurons displaying frequency tuning), characterized by increased firing within a specific frequency range and decreased firing in another. Less commonly, frequency-tuned neurons exhibited either a selective enhancement (4 neurons, 15%) or selective suppression (5 neurons, 18%) of firing within a frequency range, without a statistically significant opposing effect.

**Figure 5.**
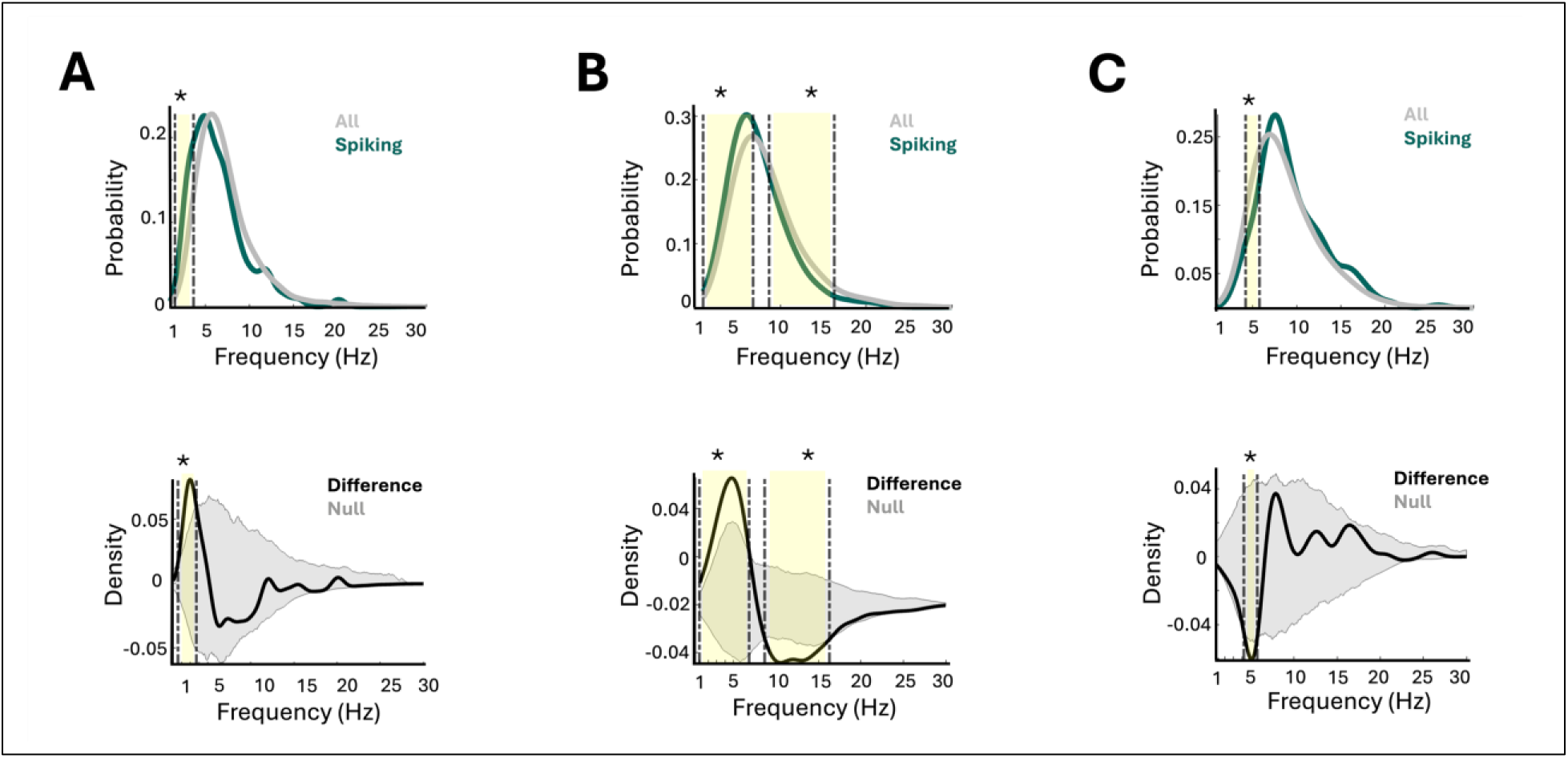
Patterns of frequency-related modulation of neuronal firing. Top panels display histograms of the normalized probability of instantaneous frequencies during spiking (pine green) and all times (gray), and the bottom panels show the difference in frequency-specific probability density between spike times and all times. The shaded gray area indicates the appropriate lower and upper percentile bounds of the null distribution generated by permutation. Asterisks (*) denote frequency bins with significant differences that survive permutation testing, also highlighted with colored shading. Three distinct patterns of frequency-related modulation are illustrated: **A.** Increased firing within a specific frequency range. **B.** Mixed modulation, with increased firing in one frequency range and decreased firing in another. **C.** Decreased firing within a specific frequency range.

To assess whether frequency preference varied across regions, we performed region-specific analyses. **Supplementary Figure 3** displays the distribution of frequencies at which maximum spiking occurred among frequency-tuned neurons, alongside the distribution of frequencies corresponding to the highest r² values across all macroelectrodes within each region. The r²-based distribution serves as an estimate of the most prevalent oscillatory frequencies in the background LFP. Across all regions, frequency tuning mostly occurred at frequencies below 10 Hz.

We next asked whether the frequency at which neurons showed increased or decreased firing was related to the prevalence of frequencies in the LFP (**Figure 6A-D**). To evaluate this, we compared the frequency associated with increased firing to the frequency corresponding to the maximum r² value from the LFP (e.g. 3Hz activity in Figure 2A) separately for each frequency tuned neuron. A signed-rank test revealed no statistically significant difference between these two measures (**Figure 6A**, p = 0.08), indicating that tuning frequencies were overlapping with the most dominant frequencies in the LFP. While not significantly different across the population, several frequency tuned neurons exhibited tuning to frequencies distant from the dominant LFP rhythm (**Figure 6A,C, Supplementary Figure 3**). Framed differently, while this overlap suggests that tuning often occurs near dominant LFP components, it does not account for the distributional shift observed in the KS analysis (see previous analysis). Instead, it suggests frequency tuning was largely confined to low frequencies and expressed with fine spectral resolution. Supporting the notion of fine-resolution tuning, we observed that the bandwidth of frequency tuning (Δf), defined as the difference between the upper and lower bounds of the tuning range, varied across neurons from 0.2 to 4.3 Hz (median = 1 Hz). The narrowest tuning (0.2 Hz) was observed within a 2.3–4.4 Hz band, while the broadest (4.3 Hz) spanned 1–23.2 Hz (**Supplementary Figure 4**). We did not observe a consistent directional relationship between the dominant LFP frequency and the frequency for increased spiking (**Figure 6C)**. These results together suggest that frequency tuning was not always driven by signal prevalence but occurred with finer spectral resolution, particularly within the low-theta range.

**Figure 6.**
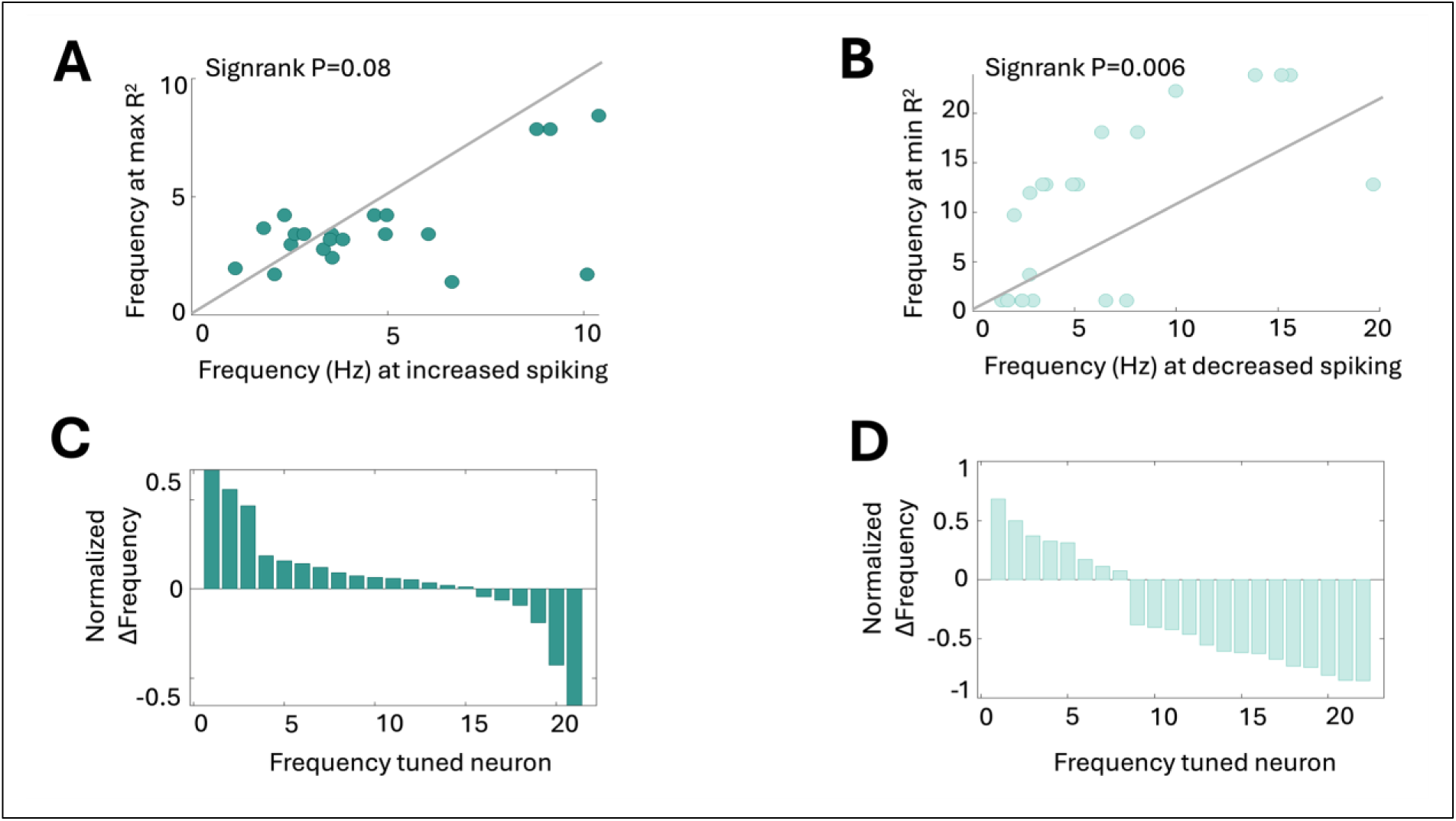
Frequency tuning reflects fine-grained preferences, not strictly tied to dominant or least prevalent LFP frequencies. **A.** Scatter plot of the frequency associated with increased spiking versus the frequency corresponding to the maximum r² from the same electrode. Data points cluster near the diagonal, indicating that while the two frequency ranges are closely aligned, they are not identical. **B.** Scatter plot of the frequency associated with decreased spiking versus the frequency corresponding to the minimum r². Points are far from the diagonal, suggesting no correspondence between the two frequency ranges. **C.** Distribution of the differences between the frequency of increased spiking and the frequency with maximum r². These frequencies span both above and below the peak of the most prevalent frequency in the LFP, highlighting fine resolution around, but not limited to, the dominant frequency. **D.** Distribution of the differences between the frequency of decreased spiking and the frequency with minimum r² from the same electrode. These frequencies also span values both above and below the least prevalent LFP frequency, again indicating that tuning does not simply reflect extremes in background spectral power.

Moreover, frequencies associated with decreased firing significantly differed from the least prevalent frequencies in the LFP (p = 0.006; **Figure 6B,D**), further supporting the idea that frequency tuning is not a byproduct of spectral density alone. In other words, the decrease in firing rates observed for some LFP frequencies was not due to a lack of LFP oscillations at those frequencies. Here again we did not observe a consistent directional relationship between the least prevalent LFP frequency and the frequency for decreased spiking (**Figure 5D**). These findings suggest that frequency tuning reflects selective modulation within specific low-frequency ranges, rather than a passive tracking of dominant or rare oscillatory components, in line with prior work (13) showing that even subtle shifts in oscillatory frequency can alter network excitability and modulate neuronal responsiveness to synaptic inputs.

Pooled over data showed that frequency tuning was most prominent at frequencies below 10 Hz (**Figure 7A**). To parallel the phase tuning analyses where we pooled phases associated with increased firing, our comparison was limited to frequencies in which tuning coincided with increased firing. A two-sample KS test revealed no significant difference between the pooled distribution of tuned frequencies and the overall frequency distribution (p = 0.44), indicating a general alignment with low-frequency activity. However, as shown in previous analyses (**Figure 6**), this overlap was not exact; tuning occurred with finer resolution, selectively targeting specific frequencies within the low-frequency range. Notably, despite observing oscillatory signals above 10Hz across all macroelectrodes (gray bars in **Figure 7A),** we did not observe frequency tuned neurons in this range (green bars in **Figure 7A**). This indicates that frequency tuning does not strictly mirror the distribution of spectral content across frequencies but instead reflects a distinct preference for lower frequencies. The pooled distribution of preferred phases, defined as the mean phase at which each phase-tuned neuron fired most, showed a tendency to cluster around the trough of the oscillation (**Figure 7B**), though this did not reach statistical significance (Rayleigh test, p = 0.13).

**Figure 7.**
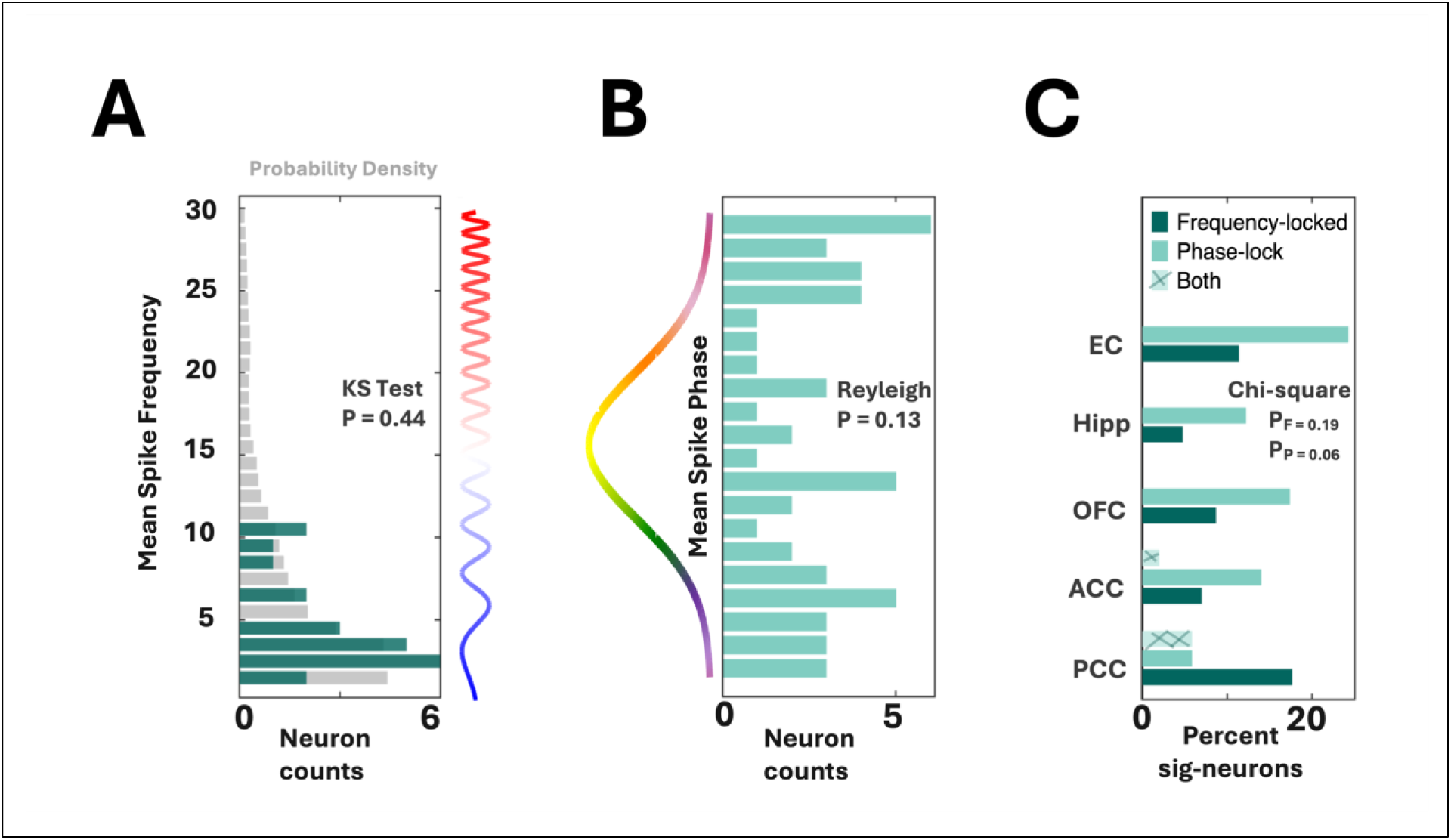
Summary of frequency and phase tuning across neurons and regions. **A.** Double axis plot showing pooled distribution of frequencies at which frequency tuning occurred across neurons recorded from all regions. The lower x-axis indicates the number of neurons (shown in pine green), while the upper x-axis shows the overall frequency probability across all macroelectrodes. Frequency tuning predominantly occurred at frequencies below 10 Hz. **B.** Pooled distribution of phases at which phase tuning occurred across neurons recorded from all regions (shown in mint green). The x-axis indicates the number of neurons. Overall, there was no consistent preference for particular phases. **C.** Regional distribution of neurons showing frequency or phase tuning, demonstrating no anatomical preference across regions.

When considering our results across different regions, we identified neurons modulated by frequency across all recorded regions: 5% in Hipp (n = 7/147), 11% in EC (n = 8/70), 7% in ACC (n = 7/100), 17% in PCC (n = 3/17), and 8% in OFC (n = 2/23). To determine whether frequency showed a region-specific distribution, we conducted a Chi-square test across Hipp, EC, and ACC. This analysis was restricted to these regions due to insufficient neuron counts in other areas, which precluded reliable statistical comparison. The regional distribution of frequency-modulated neurons was not statistically significant (Chi-square test: *X*^2^(3) = 3.26, *p* = 0.19), suggesting no regional preference for this modulation. Similarly, neurons modulated by phase were also found across all regions: 12% in Hipp (n = 17/147), 24% in EC (n = 17/70), 14% in ACC (n = 14/100), 6% in PCC (n = 1/17), and 17% in OFC (n = 4/23). The regional distribution of phase-tuned neurons did not show statistical significance (Chi-square test: *X*^2^(3) = 5.49, *p* = 0.06), indicating no clear regional preference for phase modulation either (**Figure 7C)**. The proportion of phase-tuned neurons was approximately twice that of frequency-tuned neurons in all regions, except for the posterior cingulate cortex, where frequency-tuning was more prevalent. Additionally, both phase-tuning and frequency-tuning were relatively more abundant in neurons in the entorhinal cortex compared to the other regions. Finally, only 1% of neurons (n = 3/357) were modulated by both the frequency and phase of the local LFP, suggesting that frequency-related and phase-related modulations are largely independent phenomenon. Our findings support the idea that LFP frequency and phase can independently influence neuronal excitability.

### Phase and Frequency Tuning are Independent Phenomena

Frequency and phase tuning were observed across a broad range of patients and recording sites. Frequency tuning was present in 13 out of 19 patients, while phase tuning occurred in 16 out of 19 patients. Our observations of frequency and phase tuning were not driven by a few recordings containing a high proportion of tuned neurons (e.g. different microwires in the same location); instead, modulated neurons were dispersed across many different recordings, with only a minority of neurons per microwire typically showing tuning. Among microwires with neurons showing frequency tuning, the median proportion of frequency-tuned neurons was 13% (range: 5%–100%), and the median proportion of phase-tuned neurons was 25% (range: 6.25%–50%) (**Supplementary Figure 5**). These findings suggest that frequency and phase tuning are distinct, neuron-specific phenomena rather than general properties of local field signals at individual recording sites.

We compared the firing rates of neurons exhibiting frequency tuning or phase tuning to those of untuned neurons to ensure that differences in tuning were not confounded by baseline firing rates. A Wilcoxon rank-sum test revealed no significant difference in firing rates between tuned and untuned neurons (frequency-tuned vs. untuned: p = 0.44; phase-tuned vs. untuned: p = 0.56), indicating that frequency and phase tuning were not driven by differences in overall firing rate. Firing rates were log-normally distributed (median = 1.6 Hz), consistent with previous reports (15). Across all neurons, burst indices were consistently low, and no neuron exhibited a positive burst index indicative of bursting. This suggests that the recorded neurons primarily fired in a regular, non-bursting pattern.

### Low Frequency LFP oscillatory activity drives both phase and frequency tuning

Finally, we examined the LFP spectral decomposition results obtained through ORCA to identify which spectral features were associated with phase and frequency tuning. As shown, our data-driven ORCA approach revealed variable frequency bands (**Supplementary Figure 6A**) that better capture the spectral features of frequency-tuned electrodes (**Supplementary Figure 6B**). ORCA decomposed LFP signals into 2 or 3 bands in the majority of recordings (76%). However, the number of bands was not associated with the presence of frequency or phase tuning (rank-sum test; p = 0.73 and p = 0.09, respectively). Similarly, there was no relationship between the frequency bandwidth and the presence of frequency or phase tuning (rank-sum test; p = 0.66 and p = 0.87, respectively).

To characterize regional spectral dynamics, we pooled instantaneous frequency estimates across all macroelectrodes and times and visualized their distribution probability as histograms (**Supplementary Figure 1**). Frequency distributions varied by region, with prominent peaks appearing in lower frequency ranges. For instance, hippocampal frequencies most commonly peaked around 2.3 Hz, while the ACC showed a peak near 1.4 Hz. We quantified these differences by extracting the modal frequency (i.e., the frequency bin with the highest probability) for each region. Across participants, the Hipp, EC, and ACC consistently exhibited dominant activity in the delta-theta range (1–4 Hz), while regions such as the PCC and OFC showed slightly faster dynamics. Low-frequency oscillations (<10 Hz) were prominent across all regions, accounting for 60–85% of total LFP activity in the 1–30 Hz range.

Notably, the instantaneous frequencies associated with frequency and phase tuning were predominantly in the lower range: 90% of phase or frequency tuned neurons occurred at frequencies below 10 Hz. In contrast, the full set of decomposed frequency bands across all channels extended up to 30 Hz, indicating that both frequency and phase tuning effects were largely confined to slower oscillatory activity (**Supplementary Figure 7**).

## Discussion

By analyzing single-neuron recordings from multiple brain regions in human subjects, we found that neuronal firing covaries with the instantaneous frequency of LFP oscillations, which we refer to as frequency-tuning. Frequency tuning was most prominent at frequencies below 10 Hz and was observed in each anatomical region considered (**Figure 7A**). Consistent with prior studies, we also identified neurons whose firing covaries with the phase of LFP oscillations, (i.e. phase-tuning) (**Figure 7B)**. We found that phase-tuning occurs predominantly in the lower frequency ranges, mainly between 1–10 Hz (**Supplementary Figure 7**). Furthermore, neurons showing frequency-tuning and phase-tuning were distinct. Finally, we found that phase-tuning was distributed evenly across regions (**Figure 7C**) including the Hipp, EC, ACC, PCC, and OFC, and occurred predominantly near the trough of the LFP oscillation, in line with prior studies reporting increased neural firing during trough phases. Taken together, our results show that phase-tuning and frequency-tuning occur independently of one another, suggesting that they are distinct processes likely driven by separate underlying mechanisms. In addition, they show that both phenomena occur across cingulate, frontal, and medial temporal lobe structures. These findings support the frequency-dependent modulation of neuronal firing hypothesis in humans and provide further evidence that brain oscillations influence firing patterns.

### A subset of human single neurons displays phase-tuning with a dominant role of slow oscillations

Phase modulation of neuronal spiking, often referred to as phase tuning, has been observed across species, including rodents, non-human primates, and humans (2,7,15,17,28,29). Previous studies have shown that neuronal firing synchronizes with the phase of oscillations across multiple frequency bands, including theta (30), alpha (26), beta (Canolty et al., 2012), and gamma (31). In a foundational study, Jacobs et al, provided the first evidence of phase tuning in human single-neuron recordings (2). They observed that neuronal firing was modulated by the phase of oscillations across a broad frequency range, with a stronger preference for slow rhythms, particularly within the delta and theta bands (approximately 1–10 Hz). Phase tuning did not occur at a single uniform phase but instead varied across neurons, although preferred phases tended to cluster around the trough of the oscillatory cycle. Importantly, they identified phase-tuned neurons across multiple brain regions. In line with these findings, our results also reveal phase tuning across a range of frequencies, primarily within the low-frequency range (1–10 Hz), and across various brain regions. While slightly more prominent in the entorhinal cortex, phase tuning was evident in other regions as well. Region-specific analyses confirmed that there was no single preferred phase per region; however, tuning most often occurred near the trough, consistent with prior reports of trough-locked spiking in both human and animal studies. This phenomenon aligns with theoretical frameworks such as temporal coding and communication-through-coherence, which propose that spikes occurring at specific phases, particularly near the trough, can enhance the reliability of information transfer and promote inter-regional synchrony (14,32,33).

### A subset of human single neurons displays frequency-tuning to slow oscillations independent of the phase of oscillation

Prior work in non-human primates has shown that neuronal firing can be modulated by the frequency of ongoing oscillations. Canolty et al. demonstrated that neurons in the primate frontal cortex preferentially fire during specific inter-regional phase coupling patterns at certain frequencies, forming an “internal receptive field” (12). Extending these findings to humans, studies have shown that low-frequency oscillations in the delta and theta range similarly modulate spike timing and excitability. Qasim et al. reported phase precession in human hippocampal and entorhinal neurons relative to theta rhythms, pointing to frequency-dependent modulation of spike timing (19). Additionally, Watrous et al. tested for frequency-specific phase coding in the delta band and found that the phase of delta oscillations modulates neural activity associated with different image categories (21). Finally, Roux et al. found that faster theta oscillations enhance spike-phase locking and support memory encoding (34). These studies are broadly consistent with the Spectro-Contextual Encoding and Retrieval framework which posits that frequency-specific oscillations, particularly in the theta and delta bands, organize distributed memory representations (23).

Our findings, showing prominent frequency tuning below 5 Hz across multiple brain regions, align with the Spectro-Contextual Encoding and Retrieval framework by suggesting that low-frequency oscillations play a role in modulating neuronal excitability and facilitating large-scale neural coordination. Specifically, frequency tuning below 5 Hz, a range implicated in the coordination of neural activity (35), may complement frequency-specific phase coding mechanisms involved in organizing memory representations (19,21,23). These results highlight the potential for slow oscillations to modulate neuronal excitability in a frequency-dependent manner, supporting the coordination of neural activity that underlies memory processes.

Frequency tuning, like phase tuning, was observed across multiple brain regions and phase tuning was twice as prevalent. This difference may be due to the broader frequency range associated with phase tuning. Notably, the entorhinal cortex exhibited a relative dominance in both frequency and phase tuning (**Figure 6C**), likely due to the greater prevalence of low-frequency oscillations (<10 Hz) in this region, which account for over 80% of the spectral content in the EC, compared to other brain regions (**Supplementary Figure 1**). This low-frequency environment may provide an optimal context for the emergence of both tuning types. Supporting this interpretation, Jacobs et al. emphasized the role of low-frequency oscillations in facilitating phase coding in the human brain (2). Although phase-tuned and frequency-tuned neurons show minimal overlap, suggesting they arise from distinct, independent mechanisms, both may ultimately serve the common goal of facilitating information processing.

### Regional oscillatory signatures and their relationship to frequency tuning

Previous studies have shown that brain regions have distinct oscillatory signatures as resting-state “spectral fingerprints”, like alpha rhythms in the parieto-occipital cortex and theta rhythms in the hippocampus (36). In our recordings, we observed low-frequency activity (<10 Hz) across all the regions considered, with distinct spectral peaks: ∼2 Hz in medial temporal structures (Hipp and EC), ∼1.5 Hz in the ACC, dual peaks (2 Hz and 7 Hz) in the OFC, and peaks at 2 Hz, 10 Hz, and 22 Hz in the PCC (**Supplementary Figure 1**). These regional oscillatory patterns support the idea that each area has a unique signature shaping local neural processing (37).

While neuronal frequency tuning did not exactly match the dominant LFP peaks, it occurred at nearby frequencies, suggesting that the regional oscillatory signature influences neuronal firing. For instance, a neuron may fire more at 3 Hz when the LFP peak is at 4 Hz (**Figure 3**). This shows that frequency tuning is shaped by regional oscillations, thus enabling flexible changes in firing activity with small changes in the LFP spectral distribution. This relationship aligns with EEG and TMS studies showing cortical areas have "natural frequencies" (38), driven by intrinsic resonance properties (22). Although resonance has been primarily characterized *in vitro*, our findings suggest that neurons fire more frequently when the LFP exhibits particular instantaneous frequencies, pointing to an *in vivo* correlate of intrinsic frequency preference and resonance-like dynamics.

### From frequency-sliding to frequency-tuning: manifestations of a shared oscillatory mechanism

Frequency sliding refers to the slow, moment-to-moment changes in the peak frequency of ongoing brain oscillations, reflecting the dynamic and nonstationary nature of neural activity. Unlike traditional models that treat oscillatory frequency as stable within canonical bands (e.g., alpha or theta), frequency sliding captures how the brain’s dominant rhythms shift over time in response to internal states or external inputs (13). This model proposes that instantaneous frequency fluctuations influence neuronal excitability by modulating spike thresholds. Our findings build on this framework by demonstrating that individual neurons exhibit frequency-selective spiking, preferentially firing during specific instantaneous frequencies of the LFP. This corroborates the idea that frequency sliding can shape the timing and likelihood of neuronal firing and that human single neurons are selectively responsive to certain oscillatory brain states. Cohen demonstrated lower-frequency oscillatory environments were shown to reduce spike thresholds, allowing neurons to fire in response to weaker synaptic input (13). This frequency-dependent modulation of excitability can provide a mechanistic account for our observed frequency tuning to slower oscillatory states in local field potentials.

### Non-canonical data-driven decomposition of phase and frequency of neural signal

A key distinction of our study is the use of ORCA, a data-driven algorithm that adaptively identifies instantaneous phase and frequency components from the signal itself (25), rather than imposing predefined canonical bands (e.g., delta, theta, alpha). Fixed-band approaches can force data into spectral ranges that may not be present, conflating true oscillations with broadband or aperiodic components (39) and obscuring physiological variability across individuals and species. This limitation is evident in cross-species comparisons: rodent hippocampal theta is typically ∼8 Hz, whereas human hippocampal theta is slower and spans a broader range (2,19,24,40). Data-driven approaches like ORCA, previously shown to detect non-traditional bands in human EEG and rodent recordings (25), can better capture such variability.

### Phase- and frequency-tuning in human resting-state data

Unlike many prior studies, we examined the relationship between neuronal firing and both the instantaneous phase and frequency of LFPs during the resting state. While most spike-phase coupling studies focus on task-evoked oscillations, where behavior reliably induces rhythmic activity (e.g., theta during navigation (2), memory tasks (8,15) or running (41), our analysis targeted ongoing, internally generated dynamics. Prior work has reported higher proportions of phase-locked neurons during tasks (about 70% (2) and 20% (Rutishauser et al., 2010)), likely reflecting the influence of task-driven oscillatory dynamics that enhance neuronal synchronization. In contrast, resting-state oscillations are thought to reflect intrinsic circuit properties that support large-scale coordination in the absence of external demands (13,42). Our findings highlight that even without task engagement, neuronal firing remains tuned to specific phases and frequencies, suggesting that endogenous spectral dynamics structure neuronal excitability in a behavior-independent manner. Supporting this, Cohen showed that frequency sliding itself, dynamic fluctuations in oscillatory peak frequency, was present at rest, further confirming that such spectral dynamics are an inherent feature of neural activity (13).

Together, these findings raise an important question: is frequency tuning primarily governed by internal biophysical or circuit-level properties, or do they emerge adaptively in response to behavioral context? While task-driven oscillations may transiently enhance neuronal synchrony to support cognitive functions, our results point to the possibility that such tuning mechanisms may already be scaffolded by internally generated oscillatory dynamics. Building on this, future studies can extend the present work by investigating whether frequency tuning varies as a function of behavior and if it codes for behaviorally-relevant information (i.e. “frequency coding”).

## Future directions

Our findings open several important avenues for future investigation. First, it remains to be determined whether frequency-dependent modulation of neuronal firing reflects a local circuit mechanism or a broader network-level phenomenon. While some neurons may exhibit tuning based on intrinsic dynamics or local excitability states, others may reflect coordinated activity across anatomically connected regions via shared oscillatory dynamics (6). Future work should examine whether neurons in distinct regions exhibit co-tuning to specific frequencies, potentially revealing principles of large-scale functional integration. Additionally, prior research has identified large-scale brain networks organized by frequency-specific modes of communication. A recent MEG studiy by Rosso et al. show that these networks dynamically reconfigure with sensory input, supporting the idea that oscillatory frequency acts as a scaffold for interregional coordination (43). Bridging cellular and systems levels, future research could test whether single-neuron tuning aligns with the frequency architecture of these large-scale networks, as observed in M/EEG (44), to better understand how local excitability patterns are embedded in global brain dynamics.

Second, our results raise fundamental questions of causality. Do local field oscillations actively shape neuronal excitability, or do patterns of spiking generate frequency-specific fluctuations in the LFP? Establishing the directionality of this relationship will be critical for understanding the generative mechanisms underlying frequency tuning. Rodent models offer unique advantages here, enabling cell-type-specific and circuit-level manipulations that can directly test whether altering oscillatory frequency affects spike output or vice versa.

Third, our findings emerged in the absence of overt behavioral tasks, indicating that frequency-specific modulation may reflect spontaneous network dynamics that support internally driven processing. Future studies could explore whether frequency tuning is influenced by behavioral or cognitive demands.

Finally, understanding how oscillatory frequency modulates excitability may have translational implications. The selective alignment of neuronal subpopulations to specific frequency bands suggests a novel avenue to understand the effect of neuromodulation relying on trains of electrical (45–48) or magnetic stimulation at different frequencies (49,50). By externally adjusting oscillatory frequency in a targeted brain region, it may be possible to enhance or suppress the excitability of distinct neuronal ensembles. By advancing out understanding on the coupling between neuronal activity and oscillatory dynamics, it may be possible to enable more precise, individualized neurotechnological strategies for modulating neural activity through frequency-tuned interventions, with applications in cognitive enhancement, rehabilitation, or treatment of neurological disorders.

## Limitations

Several limitations should be considered when interpreting our findings. First, the total number of recorded neurons, and the number of neurons simultaneously isolated from the same electrode contact or brain region, was limited. Future studies using high-density neural recordings, such as those enabled by Neuropixels and other modern electrode arrays (e.g., Katlowitz et al., 2025), may provide finer resolution on the population-level organization of frequency-tuned neurons.

Second, our recordings were limited to brain regions where clinical electrode implantation was necessary, which constrains the anatomical scope of our findings. It remains important to investigate frequency tuning in a wider range of cortical and subcortical areas to determine the spatial extent and specificity of this phenomenon. Animal models offer a valuable opportunity to address this limitation. In particular, testing in species that exhibit human-like low-frequency oscillatory dynamics, such as bats and non-human primates, may offer translational insight into the mechanisms and functional roles of frequency tuning. At the same time, studies in rodents can help establish the generalizability of frequency tuning across species. Together, these complementary approaches would strengthen the case that frequency-specific modulation of neuronal firing is a fundamental and conserved feature of neural coding.

Finally, our data were obtained from individuals with pharmacoresistant epilepsy. Although recordings were conducted during clinically stable periods and from epochs not suspected to reflect epileptic activity, we cannot entirely rule out the possibility that epilepsy, or its treatment, may influence oscillatory dynamics and associated neuronal responses. Future work in different neurosurgical populations without epilepsy (e.g., during intraoperative monitoring for DBS placement) and in animal models may help validate the robustness of frequency tuning across different physiological contexts.

## Conclusion

We have demonstrated that single-neuron firing in humans is modulated by the instantaneous frequency of the LFP, with many neurons showing increased or suppressed firing within distinct frequency bands. These effects occur across brain regions and reflect fine-grained, frequency-selective modulations rather than a simple tracking of the most prevalent oscillatory components. Our findings suggest that frequency tuning may provide a complementary axis to phase tuning for shaping neuronal excitability and guiding circuit-level computations.

## Methods

### Human intracranial neural recordings

We analyzed data from 19 patients (9 female) diagnosed with pharmacoresistant epilepsy who underwent intracranial monitoring at the Epilepsy Monitoring Unit (EMU) of Baylor St. Luke’s Hospital (**Supplementary Table 1**). Electrode placement was determined by the clinical team with the objective of identifying the epileptic seizure focus to assess potential surgical intervention (surgeries performed by S.S.). Electrode locations were confirmed through co-registration of pre-operative MRI with post-operative CT scans (**Figure 1A**).

Neural activity was recorded using a 512-channel Blackrock Microsystems Neuroport system. LFPs were sampled at 2 kHz with bandpass of 0.3-500 Hz, while action potentials were recorded at 30 kHz with a 300 Hz highpass filter during a 5-minute resting-state session, during which patients fixated on a cross presented on a hospital TV monitor. Line noise cancellation (60 Hz) was enabled during recording to reduce contamination from electrical mains.

Intracranial recordings were obtained using Behnke-Fried depth electrodes each equipped with a bundle of nine microwires at the tip of the probe. Eight of these microwires were designed to capture neuronal spiking activity, with insulation removed only at the tip to enable extracellular recordings. The remaining microwire was partially de-insulated (1 cm) and used as a local reference (52). LFPs were recorded from the deepest macroelectrode (**Figure 1B**).

Single-unit action potentials were isolated from the raw electrophysiological recordings using the WaveClus spike-sorting toolbox (53), and results were subsequently reviewed and curated manually. Units were considered single-units if they met several quality control criteria, including consistent spike waveform shape (e.g., amplitude, slope, and trough-to-peak duration), stable firing throughout the recording session, and minimal contamination by noise or overlapping spikes. Additional validation included visual inspection of inter-spike interval (ISI) histograms to confirm the presence of a clear refractory period, with no spikes occurring within 1 ms and fewer than 1% of ISIs falling below 3 ms. All analyses in this study focused on single-unit activity. To assess whether neurons exhibited bursting activity, we computed a burst index for each neuron based on the distribution of ISIs. The burst index was defined as the difference between the number of short ISIs (<10 ms) and long ISIs (>100 ms), normalized by their sum, yielding a value between −1 and 1. A value near +1 indicates a predominance of short ISIs suggestive of burst firing, whereas a value near −1 indicates highly regular spiking with few bursts.

This study was approved by the Medical Institutional Review Board at Baylor College of Medicine (IRB protocol number H-18112). All participants had drug-resistant epilepsy and provided informed consent for research participation.

### Signal preprocessing and rejection of epileptic activity

Data were first visually inspected to identify and exclude any clear artifactual or epileptiform activity. To minimize contamination from epileptogenic signals, data from each macroelectrode underwent an initial screening process using an automated algorithm adapted from previous studies (54). First, a fourth-order Butterworth low-pass filter was applied to remove frequencies above 80 Hz, reducing potential spike-related artifacts. Signal epochs were excluded if the envelope of the unfiltered signal exceeded five standard deviations from baseline and/or if the rectified envelope of the bandpass-filtered (25–80 Hz) signal surpassed six standard deviations (55). To ensure conservative exclusion, any ‘good data’ epoch shorter than one second was also classified as ‘bad’ and removed from analysis. This algorithm excluded approximately ∼3% of data across all LFP channels. Notably, the proportion of excluded data did not significantly differ between LFP channels containing frequency- or phase-coding neurons and those without (rank-sum test, p = 0.29, p = 0.94, respectively). Following the removal of noisy channels, the remaining data were re-referenced using the common average method.

### Electrode visualization

Intracranial electrode localization was performed using the intracranial Electrode Visualization (iELVis) software pipeline (56). Preoperative T1-weighted anatomical MRI scans and postoperative CT scans were acquired, converted to NIfTI format, and coregistered using FSL (57,58). Electrode positions were then manually identified in the aligned CT-MRI overlay using BioImage Suite (59). Electrode coordinates were then mapped to each patient’s native space using iELVis MATLAB functions (60) and visualized on the cortical surface reconstructed with Freesurfer (61). To determine anatomical placement, electrodes were assigned labels based on the most probable cortical parcellation within a 5 mm radius, following the Destrieux Cortical Atlas. For visualization of the microwires position, coordinates were extracted from the closest (deepest) macroelectrode on the Ad-Tech Behnke-Fried depth electrodes. For the purpose of LFP analysis, the deepest contact on the Ad-Tech electrode was used and visualized. Additionally, for group-level visualization, individual brain surfaces and electrode coordinates were transformed into MNI152 standard space using RAVE (62). The resulting coordinates were plotted on a the “fs average” brain template (**Figure 1A**).

### Spectral properties of neural signals

Neural oscillations are commonly examined using methods that assess activity within predefined frequency bands (e.g., Theta: 4-8 Hz). Various factors influence the occurrence and properties of band-limited neural oscillations, such as neuroanatomy, behavioral state, and the characteristics of the recording equipment (10). Moreover, band-limited activity is frequently assumed rather than explicitly measured, and the criteria for defining frequency bands differ significantly across studies. Therefore, a tool that captures oscillatory variability in neural signals beyond using canonical, fixed-frequency bands is needed to measure neural oscillations with more precision and less researcher bias in frequency band identification. To accomplish that, we used previously developed the Oscillatory ReConstruction Algorithm (ORCA) by our group (25) to analyze the spectral properties (instantaneous frequencies and phases) of neural signals within adaptively determined frequency bands. Generally, ORCA is an unsupervised approach that integrates two novel methods for identifying frequency bands. It utilizes the instantaneous amplitude, phase, and frequency of each band to reconstruct neural signals and evaluate spectral decomposition accuracy through four distinct models by selecting spectral estimates based on the best-performing model. This approach minimizes the need for extensive hyperparameter tuning. From these adaptively filtered signals, ORCA derives instantaneous phase and frequency estimates, while excluding outlier values arising from phase slips by masking them as NaNs. ORCA thus enables high-resolution tracking of dynamic spectral components without imposing rigid *a priori* frequency constraints.

From a signal processing perspective, ORCA segments the broadband signal using multiple strategies, estimates spectral features (amplitude, phase, and frequency) in each band, and reconstructs the signal from these estimates. Notably, as in our prior work (25), we estimated the prevalence of oscillations at each frequency by quantifying the fit between the LFP and a reconstructed signal based on activity at each point frequency (e.g. **Figure 2A**). The maximum and minimum values were then extracted for each LFP to generate results for Figure 6. Overall decomposition accuracy (i.e. goodness-of-fit) is assessed by the coefficient of determination (r²) between the original and reconstructed signals using activity across all frequencies, allowing selection of the most representative band structure.

### Statistical analyses

For this analysis, we focused on signals within the 1–30 Hz range to minimize the influence of higher-frequency spike-related artifacts, which can confound the interpretation of LFP signals (10,31,63). To strictly limit the analysis to the 1–30 Hz range, we replaced all time points with NaN wherever the instantaneous frequency exceeded 30 Hz, thereby excluding high-frequency components from all subsequent analyses. Our primary goal was to investigate how neuronal firing preferences vary based on the frequency and phase of ongoing oscillatory activity. To examine this, we compared the distribution of band-specific instantaneous frequencies recorded from the deepest macroelectrode (i.e., the closest to the microwires) across all times with the distribution at time points where spiking occurred. A two-way Kolmogorov-Smirnov (KS) test was applied to assess differences between these distributions. To account for the possibility of chance-level effects, we implemented a permutation procedure by performing the KS test 1000 times per neuron using a shuffled version of the instantaneous frequency data. This shuffling was done using a circular shift (circshift in MATLAB), preserving the temporal structure of each session. The resulting surrogate distribution of KS test statistics was used to determine statistical significance, with p-values computed by ranking the observed test statistic relative to the surrogate distribution **(Figure 1C)**.

For cases where the KS test revealed a significant difference, we identified the specific frequency range driving this effect by computing kernel density estimates (KDEs) of the instantaneous frequency distributions at spike times and all times. We then quantified their absolute difference across a common frequency grid to isolate the frequency ranges contributing most to the observed divergence. To assess the statistical significance of this frequency-specific difference, we used a permutation-based procedure with circular time-shifting (as described above). For each permutation, we repeated the KDE comparison and generated a null distribution of differences. The observed difference was then compared against the null distribution. Statistical significance was defined as values falling outside the α-corrected thresholds of the null distribution. This interval delineated the frequency range that contributed most strongly to the observed tuning effect. By examining differences in spiking activity within this range, we were able to identify whether neuronal firing was significantly enhanced or suppressed, thereby revealing both the directionality and selectivity of frequency tuning for each neuron **(Figure 5)**.

To assess phase-locked neural firing, we used the Rayleigh test, extracting the LFP phase at each spike occurrence within detected frequency bands. The mean phase angle was computed to calculate the central tendency of phase values. Circular statistical analyses were performed using the CircStat2012a MATLAB toolbox.

All analyses were conducted separately for each frequency band, and multiple comparisons were addressed using Bonferroni correction for the KS and Rayleigh tests across ORCA-detected frequency bands. The threshold for statistical significance was set at p < 0.05, with corrections applied where necessary. All the analyses were carried out in in MATLAB (v2024b, MathWorks, MA, USA)

## Acknowledgments

We thank our patients and their families for taking part in this research. We are grateful to the hospital staff for their assistance in keeping patients comfortable and cared for throughout their stay at the hospital.

## CRediT

**Zahra Jourahmad –** conceptualization, investigation, data curation, formal analysis, software, validation, visualization, writing - original draft

**Raissa K. Mathura –** investigation, writing - review and editing

**Layth S. Mattar –** investigation, writing - review and editing

**Melissa C. Franch –** investigation, writing - review and editing

**Danika L. Paulo –** investigation, writing - review and editing

**Mohammed Hasen –** investigation, writing - review and editing

**Nicole R. Provenza –** resources, writing - review and editing

**Ben Y. Hayden –** resources, writing - review and editing

**Sameer A. Sheth –** resources, funding acquisition, project administration, writing - review and editing

**Eleonora Bartoli –** conceptualization, project administration, investigation, supervision, writing - review and editing

**Andrew J. Watrous –** conceptualization, project administration, methodology, investigation, supervision, software, writing - review and editing

