## Supplementary material for "Human neuronal firing is modulated by the frequency of local field potential oscillations"

**Supplementary Table 1.** Patients' demographics

| **Gender** | | **Race** | | **Age (years)** | |
|---|---|---|---|---|---|
| Female | 9 | White | 11 | 20-30 | 7 |
| Male | 10 | Black | 3 | 31-40 | 3 |
| | | Hispanic | 2 | 41-50 | 6 |
| | | Asian | 2 | 51-60 | 3 |
| | | Multi-race | 1 | | |

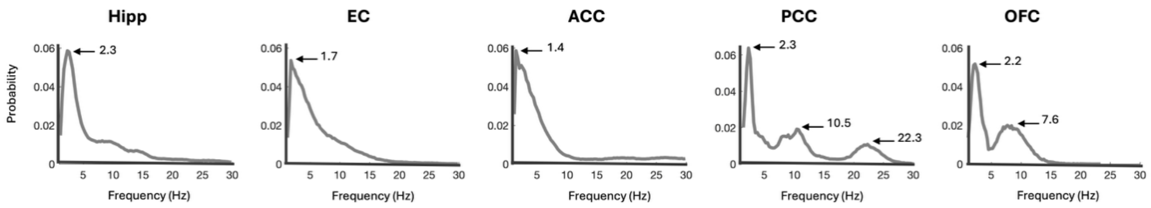


**Supplementary Figure 1.** The figure shows the probability density of instantaneous frequencies filtered between 1 and 30 Hz for each region. All regions exhibit a dominant peak in the low-theta range (<4 Hz). Additionally, PCC and OFC signals display secondary peaks. Hipp peak at ~2.3 Hz; 73% of the frequency content below 30 Hz falls under 10 Hz. EC peak at ~1.7 Hz; 80% of frequencies <30 Hz are <10 Hz. ACC peak at ~1.4 Hz; 75% of frequencies <30 Hz are <10 Hz. PCC peaks at ~2.3 Hz, 10.5 and ~9 Hz; 60% of frequencies <30 Hz are <10 Hz. OFC peaks at ~2.2 Hz and ~7.6 Hz; 84% of frequencies <30 Hz are <10 Hz.

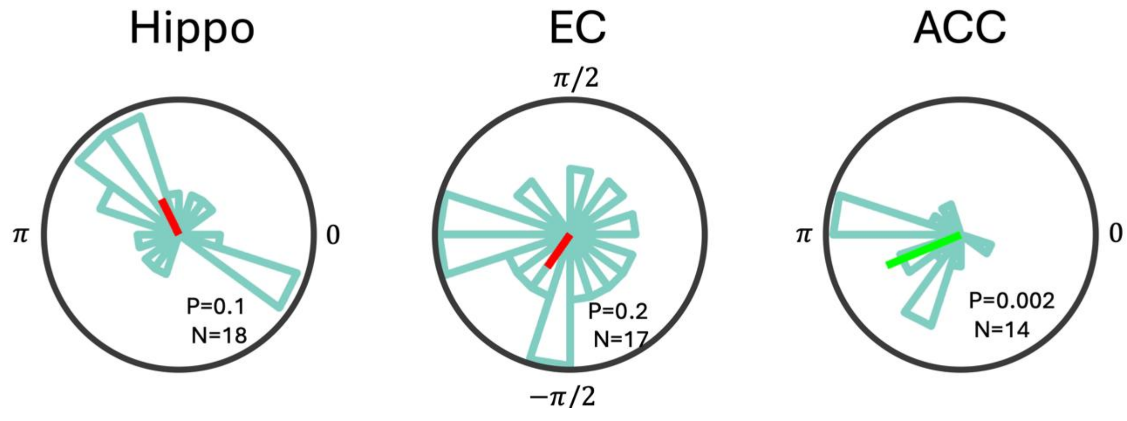


**Supplementary Figure 2.** Pooled phase distributions of neurons showing significant phase tuning, separated by region. Rayleigh tests revealed no significant phase preference in pooled phase distributions of hippocampal and entorhinal neurons, as indicated by their mean phase vectors (red). In contrast, ACC neurons exhibited a significant preference for the trough of the oscillation, as indicated by the mean phase vector (green).

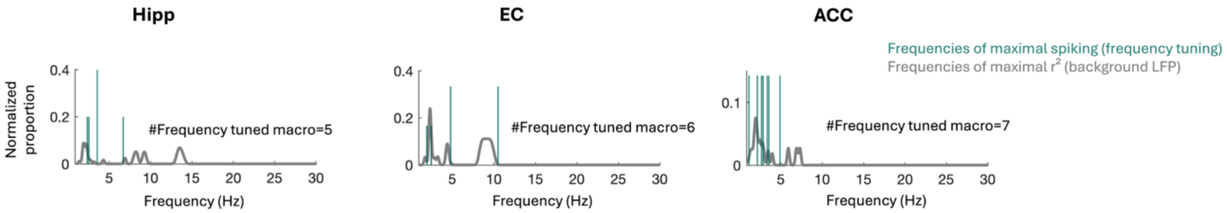


**Supplementary Figure 3.** Region-specific frequency preferences of frequency-tuned neurons compared to background LFP activity recorded from specific regions. For each region, the pine green bars show the distribution of frequencies at which frequency-tuned neurons exhibited maximal spiking activity. Overlaid in gray is the distribution of frequencies corresponding to the highest $r^2$ values across all macroelectrodes in that region, representing the most prevalent background oscillatory frequencies. Across regions, frequency tuning predominantly occurred below 5 Hz and did not consistently align with the dominant peaks of the background LFP distribution.

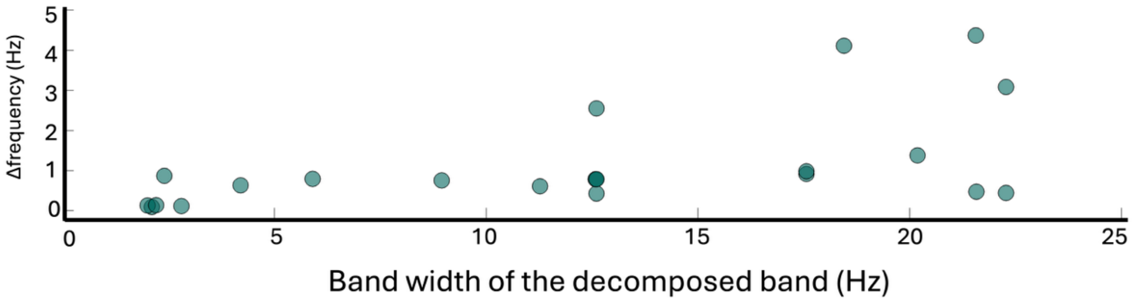


**Supplementary Figure 4.** Relationship between frequency-tuning resolution and oscillatory-band bandwidth. The x-axis shows the bandwidth of each ORCA-decomposed band (upper frequency − lower frequency), while the y-axis plots Δfrequency: the absolute difference between the frequency at which significant tuning is observed and the frequency that maximizes the model fit (peak $r^2$). The narrow spread of Δfrequency values across a range of bandwidths indicates fine resolution in the frequency tuning process.

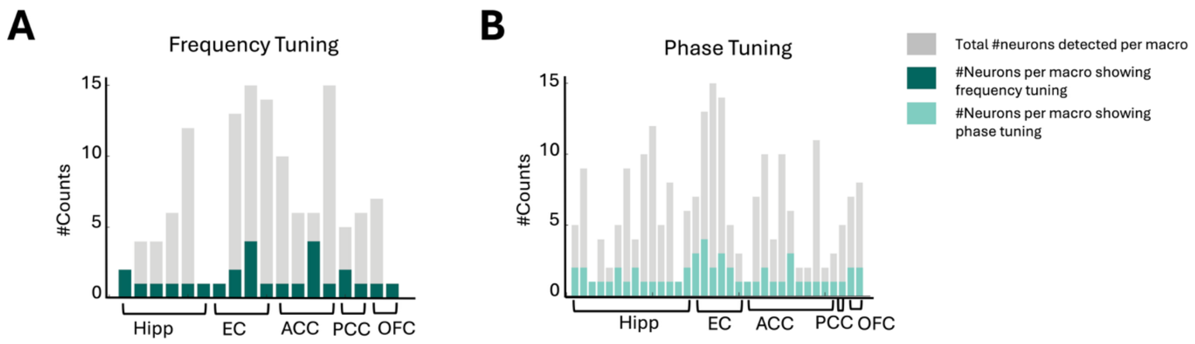


**Supplementary Figure 5.** Proportion of frequency- and phase-tuned neurons per macroelectrode across brain regions. **A.** Proportion of single neurons exhibiting frequency tuning, calculated relative to the total number of isolated single neurons from microwires stemming from the tip of the same macroelectrode. Data are shown across regions for 18 macroelectrodes from which at least one corresponding single neuron exhibited frequency tuning. **B.** Proportion of single neurons exhibiting phase tuning, relative to the total number of isolated single neurons from microwires stemming from the tip of the same macroelectrode. Data are shown across regions for 34 macroelectrodes from which at le

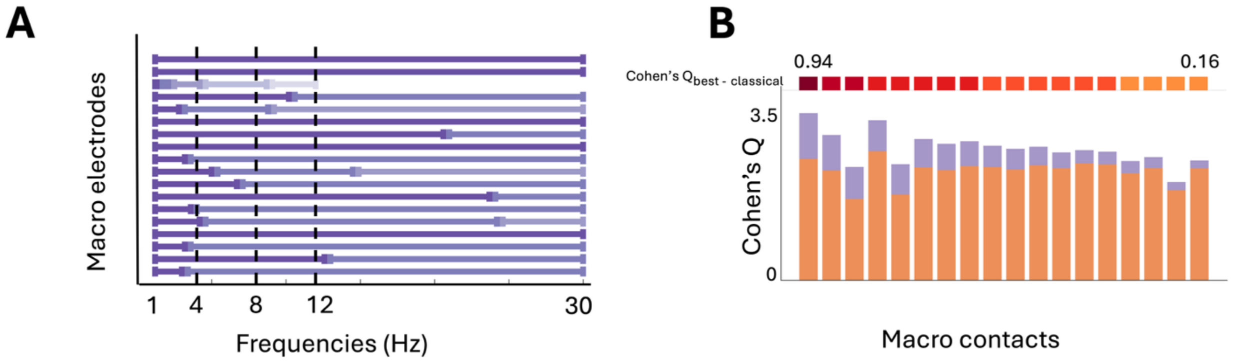


**Supplementary Figure 6. A.** Frequency tuning results for 18 macro contacts where single neurons exhibited significant tuning to their LFP. Channels are sorted by descending Cohen's Q difference, based on ORCA-derived decomposition with a variable number of bands. **B.** Comparison of reconstruction quality between ORCA and classical methods across channels, showing the improvement in fit achieved by ORCA, quantified by Cohen's Q difference. The color scheme matches that of Figure 2, with **purple** indicating the ORCA-selected method and **orange** denoting the classical method.

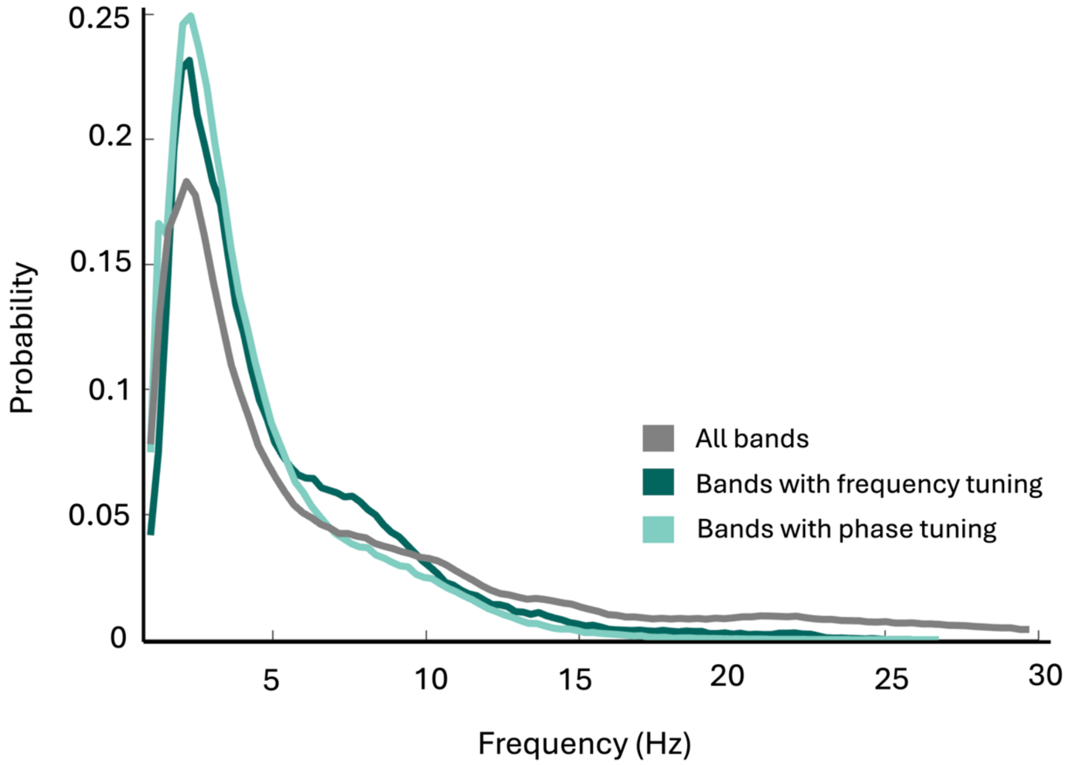


**Supplementary Figure 7.** The figure compares the distributions of instantaneous frequencies at which phase tuning and frequency tuning occurred to the distribution of frequencies across all time points, regardless of tuning. Both phase and frequency tuning preferentially occur at low theta frequencies, resulting in tuning-specific distributions that differ markedly from the overall frequency distribution.
